# Spatial organization of AQP4 channels in the human brain: links with perfusion, edema, and disease vulnerability

**DOI:** 10.64898/2026.02.18.706679

**Authors:** Tahmineh Taheri, Asa Farahani, Zhen-Qi Liu, Eric G. Ceballos, Adil Harroud, Alain Dagher, Bratislav Misic

**Affiliations:** Montréal Neurological Institute, McGill University, Montréal, Québec, Canada

## Abstract

Aquaporin-4 (AQP4) water channels support the glymphatic system, a brain-wide pathway that clears cerebral waste products. Despite its importance, the whole-brain organization of this system in humans remains underexplored. Here we use AQP4 gene expression as a molecular anchor to reconstruct a whole-brain glymphatic-related topography and link it to vascular physiology, edema, and neurodegenerative vulnerability. We find that AQP4 expression is highly organized across the brain, peaking in subcortical, ventral, and periventricular territories, consistent with a clearance axis near cerebrospinal fluid reservoirs and perivascular interfaces. Linking AQP4 expression to vascular organization, we find that AQP4-enriched regions show lower normative blood perfusion and lower vein density, suggesting that this axis is not simply explained by vascular supply or large-vessel anatomy. Turning to neurodegeneration, we find that atrophy patterns across multiple neurodegenerative diseases co-localize with AQP4 expression—most strongly for tau and TDP-43 proteinopathies—and high-atrophy regions lie close to AQP4 hotspots in both anatomical space and structural connectome space. Furthermore, incorporating normative PET markers of neuroinflammation typically strengthens the spatial alignment between AQP4 expression and disease atrophy, suggesting that inflammatory tone and glymphatic-related architecture jointly shape vulnerability. Finally, we show that peritumoral edema is most frequent in AQP4-enriched regions and is further shaped by regional inflammatory tone. Collectively, this work highlights a whole-brain glymphatic organization that relates to diverse aspects of brain physiology and vulnerability.

## INTRODUCTION

The brain is organized across multiple spatial scales [11]. Genomic gradients [14, 45] shape the spatial arrangement of cell types and laminar differentiation [35, 66, 79, 131], which in turn determine the wiring of neural circuits and the emergence of patterned neural activity [9, 12, 31, 100, 120, 128]. This complex system is perpetually active, requiring constant exchange of fluids to deliver resources (such as glucose and oxygen) and clear waste products (such as metabolites and proteins) [98]. While the former is relatively well studied, including metabolism and blood flow [19, 108], the latter—known as the glymphatic system—[55] is comparatively less understood.

Central to the glymphatic system are aquaporin-4 (AQP4) integral membrane channel proteins [30, 50, 80]. AQP4 channels are located primarily on astrocyte endfeet that line perivascular spaces, supporting the exchange of cerebrospinal fluid (CSF) and interstitial fluid (ISF) [84]. The glymphatic system, powered by AQP4 water channels, drives bulk transport of CSF from the subarachnoid space along periarterial spaces; after mixing with interstitial fluid in the parenchyma, it exits the parenchyma through perivenous spaces [38, 50, 61]. This pro-cess regulates directional interstitial fluid movement, waste clearance, and brain health. Despite its importance, how this system is organized at the whole-brain level is unknown.

The spatial arrangement of AQP4 channels is an important question, as multiple neurological diseases are associated with AQP4 dysfunction [93, 97]. Neuromyelitis optica spectrum disorder (NMOSD) is an autoimmune disease in which AQP4 antibodies bind to the channel, causing inflammation and disrupting AQP4 function [46, 54, 67]. NMOSD brain lesions show preferential involvement of regions with high AQP4 expression [1]. Beyond NMOSD, AQP4 dysfunction is also implicated in neurodegenerative diseases. The loss of perivascular AQP4 polarization accompanies aging and neurodegeneration, with potential consequences for interstitial solute clearance and the accumulation of misfolded proteins such as amyloid-*β*, tau, and TDP-43 [49, 53, 65, 92, 133]. In parallel, impaired AQP4-mediated water transport can disturb fluid homeostasis and contribute to vasogenic edema, as excess interstitial fluid is not efficiently cleared from brain tissue [62, 75, 89]. Neuroinflammation may exacerbate these processes by altering astrocyte reactivity and further disrupting perivascular AQP4 polarization, creating a cycle in which inflammation, glymphatic dysfunction, impaired waste clearance, and neurodegeneration reinforce each other [15, 30, 47]. Together, AQP4 supports healthy brain waste clearance and water balance, and its dysfunc-tion is linked with neurodegeneration and edema.

Here we study the whole-brain organization of AQP4. We first map the expression of AQP4 gene in cortex and subcortex and use this map as a molecular anchor for the glymphatic system. Looking at healthy brain function, we ask how the spatial distribution of AQP4 relates to regional differences in blood perfusion and vein density. Looking at dysfunction, we investigate how the spatial distribution of AQP4 relates to regional differences in vulnerability to neurodegenerative disease and edema. We also ask whether normative PET markers of neuroinflammation improve these associations. Finally, we assess the extent to which these results are specific to AQP4 by repeating key analyses for related aquaporins, AQP1 and AQP9.

## RESULTS

### Mapping AQP4 in the brain

AQP4 monomers are encoded by the AQP4 gene. AQP4 expression is estimated across the brain using microarray gene expression data from the Allen Human Brain Atlas (AHBA) [45]. Quality control includes intensity-based filtering and differential stability analysis [76] (see *Methods*). Expression values are parcellated to the Schaefer 400 cortical [107] and Melbourne subocortical S4 [117] atlases, yielding a whole-brain parcellation of 454 regions. To validate the spatial distribution of AHBA microarray measurements, we compare it with RNA-seq data from AHBA (Fig. S1a) and the Human Protein Atlas (HPA) as an independent dataset (Fig. S1b) [111, 119]. In addition, we compare AQP4 expression with an *in vivo* tracer assay of glymphatic transport. We use regional brain enrichment of an intrathecally administered MRI tracer (gadobutrol) measured 24 hours after administration in a previously reported reference cohort [101] (see *Methods*). AQP4 expression shows spatial correspondence with the 24 hour tracer enrichment map (Fig. S2), supporting AQP4 topography as a molecular proxy for glymphatic organization in humans.

Figure 1a shows the spatial pattern of AQP4 expression across the brain. AQP4 is expressed throughout the brain, with greater expression near subcortical, periventricular, and ventral regions. This periventricular enrichment is also evident in a volume rendering of high AQP4 expression values (Fig. S3). Within the subcortex, AQP4 expression is highest in amygdala and basal ganglia territories, including pallidum, caudate, putamen, and nucleus accumbens, and is also elevated in ventral thalamic nuclei. In contrast, hippocampal regions show lower expression overall, with the lowest values in posterior subdivisions (Fig. 1b). In the cortex, we find greater AQP4 expression throughout the limbic and transmodal cortices, including cingulate cortex, insula, medial temporal lobe, and medial prefrontal cortex. Conversely, AQP4 expression is lowest in unimodal cortex, such as primary visual, somatosensory, and auditory cortex (Fig. 1c). In sum, the expression of AQP4 is highly organized across the brain. In the following sections, we investigate the extent to which this organization aligns with other related systems [40].

**Figure 1.**
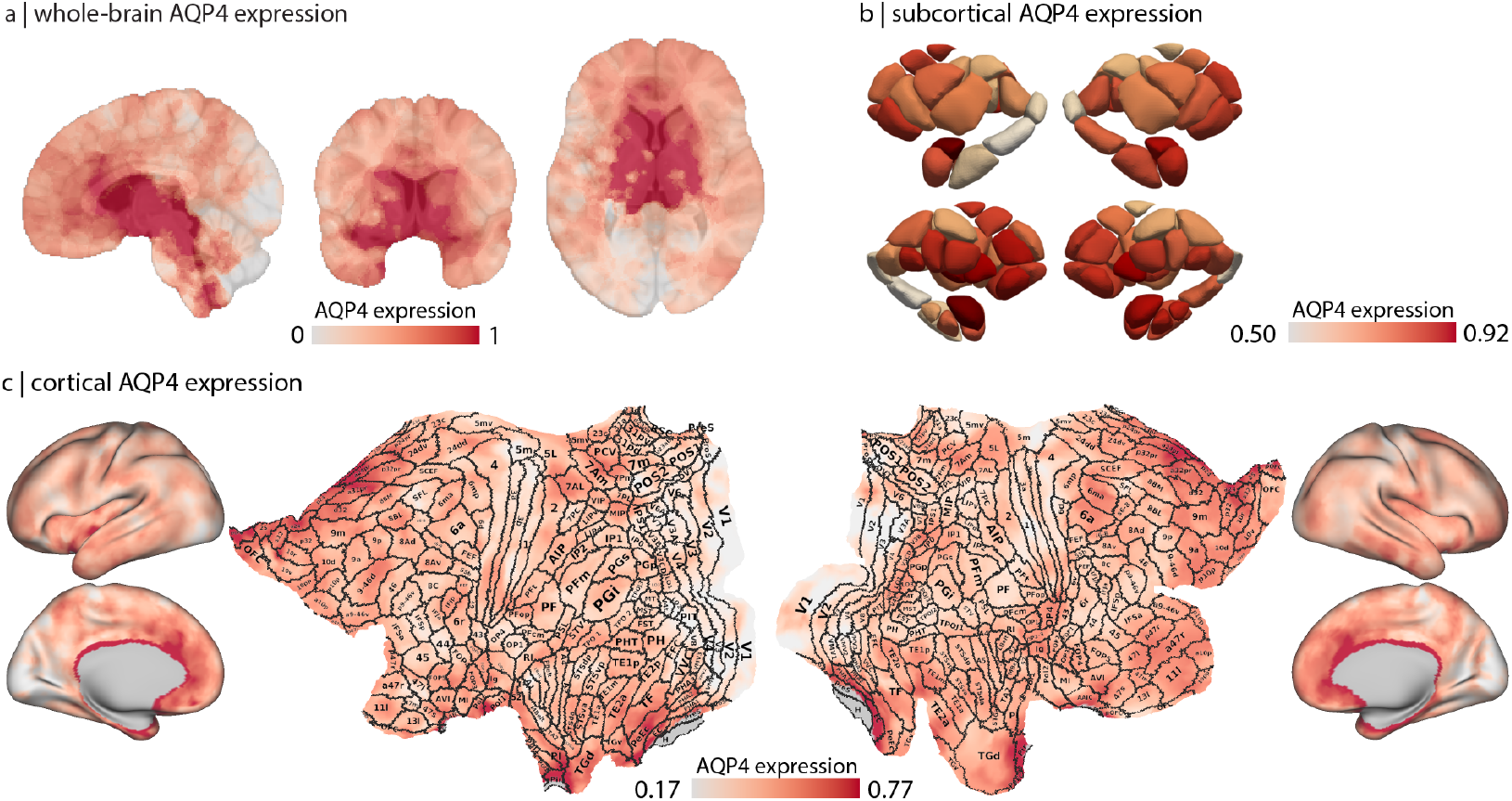
Whole-brain, cortical, and subcortical AQP4 expression maps. Donor-averaged AQP4 expression from the Allen Human Brain Atlas (AHBA) [45] processed with the abagen workflow (see *Methods*) [76]. (a) A dense, interpolated volumetric AQP4 expression map is displayed on a T1-weighted MNI template in sagittal, coronal, and axial views (MNI152-NonLinear2009cAsym, 1 mm; slice indices: sagittal=102, coronal=137, axial=81). (b) Subcortical AQP4 expression parcellated to the Melbourne subcortical S4 atlas [117], shown as 3D parcel surfaces across multiple views. Subcortical parcel names and abbreviations are listed in Table S1; labeled renderings are shown in Fig. S4. (c) Cortical AQP4 expression is shown on fsLR inflated and flat surfaces by projecting the dense volumetric map to the cortical surface; borders and areal labels from the multi-modal Glasser parcellation are overlaid on the flat surface for anatomical reference [33]; the full Glasser nomenclature is provided in Table S2.

### AQP4 channels and vascular organization

Given the molecular and cellular links between glymphatic transport and the cerebrovascular system, we first ask whether AQP4 expression co-varies with regional blood perfusion. To address this question, we estimate cerebral blood perfusion using pseudo-continuous arterial spin labeling (ASL) magnetic resonance imaging (MRI) and derive a normative perfusion map in the HCP Lifespan dataset (1305 subjects; Fig. 2a,b; see *Methods*) [13, 23, 41, 112]. We find that AQP4 expression and blood perfusion are significantly anticorrelated (Fig. 2e, top; *r* = −0.588, *p*_MSR_ = 4 ×10^−4^), indicating that the two systems are organized along opposing spatial gradients.

**Figure 2.**
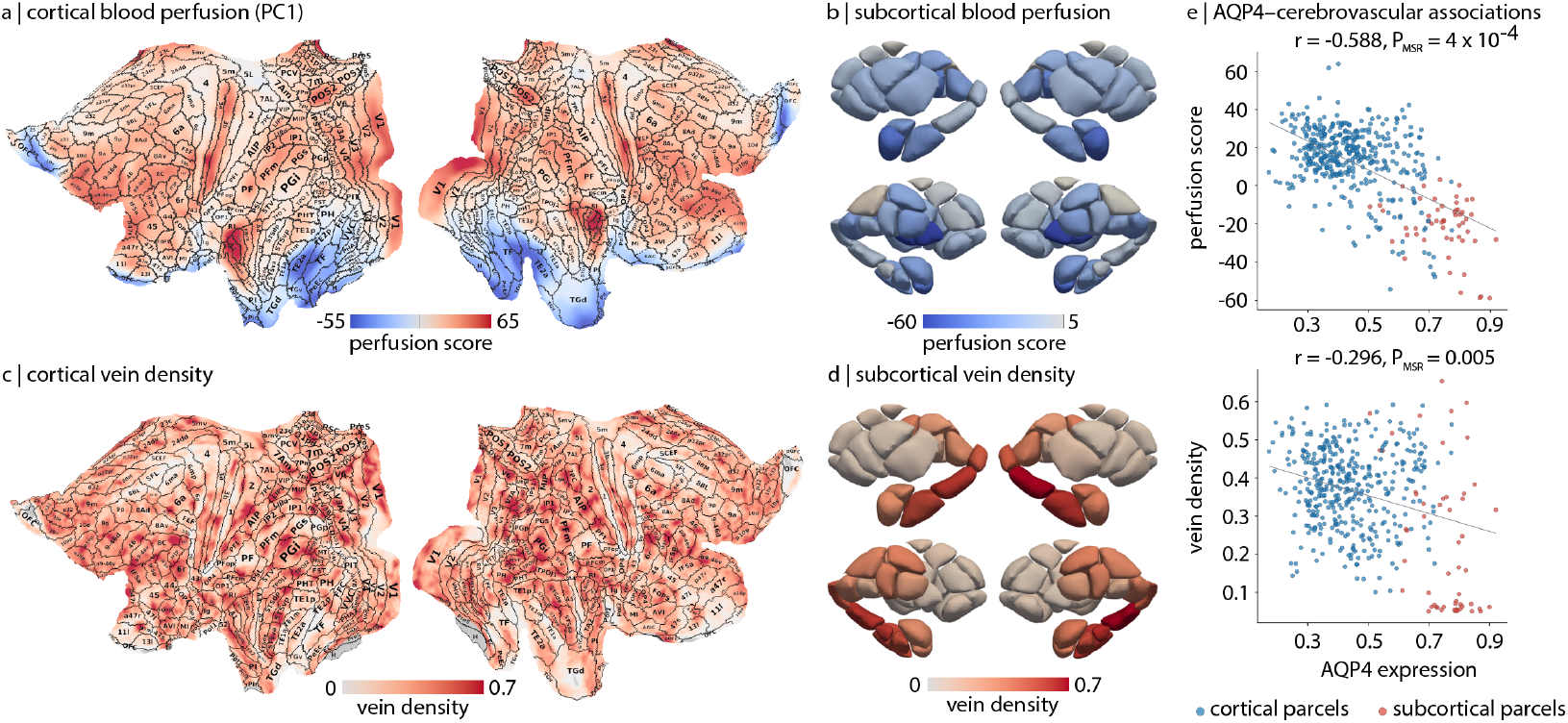
Spatial association between AQP4 expression and vascular organization. (a) Cortical cerebral blood perfusion (PC1), derived from the HCP Lifespan dataset [13, 23, 41, 112], is shown on a 2D flat fsLR surface. Borders and areal names from the multi-modal Glasser parcellation are overlaid on the flat surface [33]; the full Glasser nomenclature is provided in Table S2. (b) Subcortical blood perfusion parcellated to the Melbourne subcortical S4 atlas [117] and shown as 3D parcel surfaces across multiple views. Subcortical parcel names and abbreviations are listed in Table S1, and labeled parcel renderings are shown in Fig. S4. (c,d) Cortical and subcortical maps of vein density from the VENAT atlas [48], shown in the same space as panels (a,b). (e) Pearson correlations between AQP4 expression from the Allen Human Brain Atlas [45] (x-axis) and vascular measures (y-axis). Top: correlation with blood perfusion (*r* = −0.588, *p*_MSR_ = 4 × 10^−4^). Bottom: correlation with vein density (*r* = 0.296, *p*_MSR_ = −0.005). Each point is a parcel (blue: Schaefer 400 cortical parcels [107]; red: Melbourne S4 subcortical parcels [117]). Grey lines show least-squares fits. *p*_MSR_ values are computed using Moran spectral randomization (*N*_MSR_ = 10 000) [125] (see *Methods*).

Next, we examine whether AQP4 aligns with venous anatomy. To do so, we estimate regional differences in vein density using the Venous Neuroanatomy (VENAT) atlas, which is derived from high-resolution 7 T quantitative susceptibility mapping (QSM) in healthy adults (20 subjects; Fig. 2c,d; see *Methods*) [48]. We find that AQP4 expression is anticorrelated with vein density (Fig. 2e,bottom; *r* = −0.296, *p*_MSR_ = 0.005). This association is driven primarily by subcortical parcels: deep nuclei with high AQP4 expression tend to reside in territories with low VENAT-derived vein density, whereas cortical parcels show little systematic relationship. Because VENAT is most sensitive to larger draining veins, these findings suggest that AQP4-rich glymphatic territories are not concentrated around macro–venous trunks, consistent with a greater role for microvascular and perivascular pathways. Together, these results suggest that the spatial distribution of AQP4 is not explained simply by regional blood flow or large-vein density, pointing to a partially distinct glymphatic-related axis in the human brain.

### AQP4 channels and susceptibility to neurodegeneration

Neurodegenerative diseases are characterized by the accumulation of misfolded, aggregate-prone proteins in the brain, including amyloid-*β*, tau, and TDP-43 [103, 113]. Because these proteins are cleared in part through perivascular fluid transport, regional differences in glymphatic outflow may contribute to where pathology occurs. One possibility is that regions with high AQP4 expression lie along bottleneck pathways for glymphatic outflow, where interstitial solutes and proteins are more likely to accumulate over time, increasing susceptibility to atrophy. Alternatively, vulnerability increases when perivascular clearance is disrupted. In aging and neurodegenerative disease, AQP4 is frequently mislocalized, with reduced perivascular polarization at astrocyte endfeet. Such changes can impair CSF– ISF exchange and slow the clearance of interstitial solutes [43, 49, 53, 97, 133]. Slower clearance is expected to increase the residence time of soluble proteins in the extracellular space, facilitating protein accumulation and neurodegeneration. Together, these considerations motivate the hypothesis that brain regions with higher AQP4 expression may be preferentially vulnerable to atrophy, whether because they sit along key outflow bottlenecks or because they depend more strongly on intact AQP4-supported perivascular clearance.

To test this idea at the whole-brain level, we ask whether the spatial distribution of AQP4 relates to patterns of grey matter loss across neurodegenerative diseases. We use voxel-based morphometry maps from a pathologically confirmed dementia cohort [42] comprising early-onset Alzheimer’s disease (EOAD), late-onset Alzheimer’s disease (LOAD), presenilin 1 mutation carriers (PS1), three-repeat tauopathy (3Rtau), four-repeat tauopathy (4Rtau), frontotemporal lobar degeneration with TDP-43 type A pathology (TDP-43A), frontotempo-ral lobar degeneration with TDP-43 type C pathology (TDP-43C), and dementia with Lewy bodies (DLB) (see *Methods*). Atrophy is estimated using voxel-based morphometry (VBM) applied to the T1-weighted images, which is then parcellated to the same Schaefer–Melbourne atlas as the AQP4 map [107, 117]. We then correlate AQP4 expression with each atrophy pattern and assess significance using Moran spectral randomization to account for spatial autocorrelation [78, 123, 125] (see *Methods*).

Across diseases, we observe differences in how strongly atrophy co-localizes with AQP4 expression (Fig. 3). TDP-43 and tau proteinopathies with similar patterns of frontotemporal volume loss show the strongest positive correlations with AQP4 expression, including TDP-43C (*r* = 0.541; FDR-corrected, *p*_MSR_ = 0.017), TDP-43A (*r* = 0.436; FDR-corrected, *p*_MSR_ = 0.037), 4Rtau (*r* = 0.556; FDR-corrected, *p*_MSR_ = 0.017), and 3Rtau (*r* = 0.600; FDR-corrected, *p*_MSR_ = 0.045). These pathologies are characterized by marked atrophy of frontal, insular, and anterior temporal cortices, regions in which AQP4 expression is also relatively high. More modest but still significant associations are observed for LOAD (*r* = 0.257; FDR-corrected, *p*_MSR_ = 0.017) and DLB (*r* = 0.262; FDR-corrected, *p*_MSR_ = 0.017), whose atrophy patterns emphasize medial temporal and temporoparietal regions. By contrast, EOAD and PS1 mutation carrier groups show weak and non-significant correlations with AQP4 expression (both |*r*| < 0.1; FDR-corrected, *p*_MSR_ *>* 0.4), consistent with their more extensive temporoparietal and sensorimotor volume loss, which only partially overlaps with the AQP4 gradient. These results suggest that brain regions with high AQP4 expression are more susceptible to neurodegeneration.

**Figure 3.**
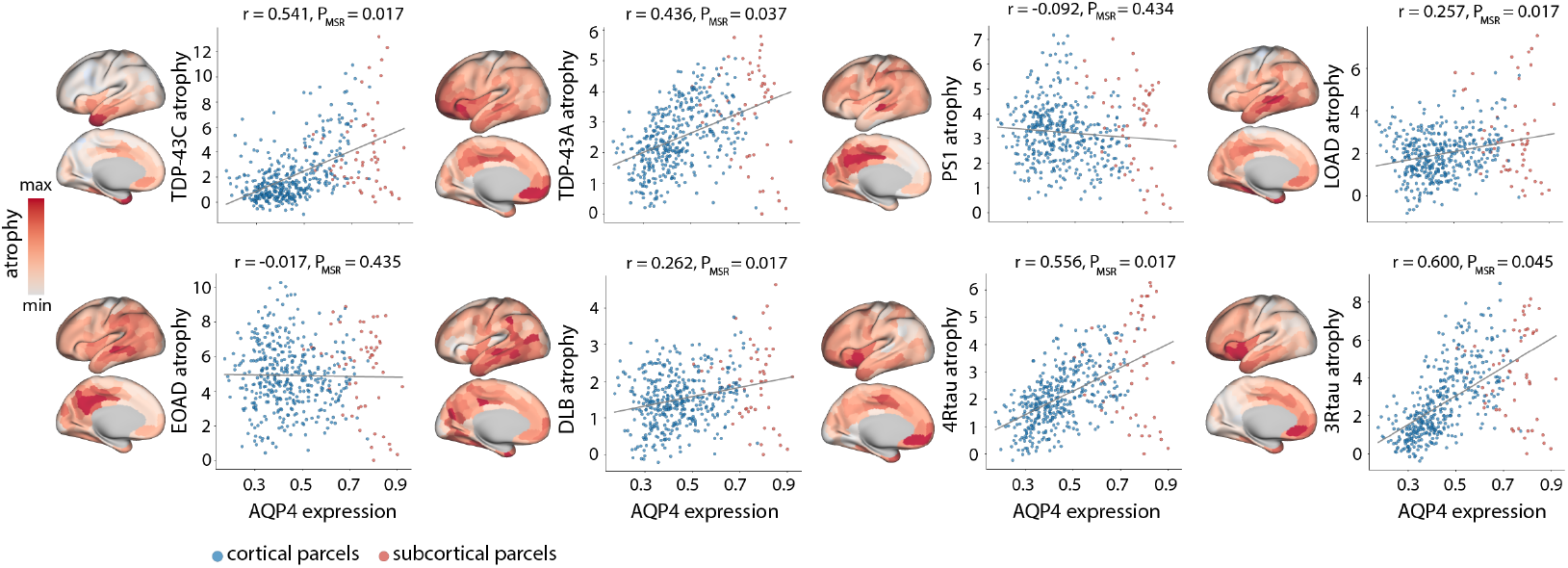
Co-localization of AQP4 expression with dementia atrophy patterns. Insets show cortical atrophy maps on left hemisphere fsLR surfaces. Scatterplots show parcel-wise Pearson correlations between AQP4 expression (x-axis) and regional atrophy (y-axis) from a pathologically confirmed dementia cohort [42]. Panels (left-to-right) are: frontotemporal lobar degeneration with TDP-43 type C pathology (TDP-43C; *r* = 0.541; FDR-corrected, *p*_MSR_ = 0.017), frontotemporal lobar degeneration with TDP-43 type A pathology (TDP-43A; *r* = 0.436; FDR-corrected, *p*_MSR_ = 0.037), presenilin 1 mutation carriers (PS1; *r* = −0.092; FDR-corrected, *p*_MSR_ = 0.434), late-onset Alzheimer’s disease (LOAD; *r* = 0.257; FDR-corrected, *p*_MSR_ = 0.017), early-onset Alzheimer’s disease (EOAD; *r* = −0.017; FDR-corrected, *p*_MSR_ = 0.435), dementia with Lewy bodies (DLB; *r* = 0.262; FDR-corrected, *p*_MSR_ = 0.017), four-repeat tauopathy (4Rtau; *r* = 0.556; FDR-corrected, *p*_MSR_ = 0.017), and three-repeat tauopathy (3Rtau; *r* = 0.600; FDR-corrected, *p*_MSR_ = 0.045). Each point is a parcel (blue: cortical parcels; red: subcortical parcels). Grey lines show least-squares fits. *p*_MSR_ values are computed using Moran spectral randomization (*N*_MSR_ = 10 000).

### Proximity of AQP4 hotspots to high-atrophy regions

So far, we have shown that AQP4 expression is associated with disease-specific atrophy patterns across multiple neurodegenerative diseases. Here, we ask whether the most affected regions lie proximal to AQP4-enriched territories. To address this question, we define AQP4 hotspots as parcels in the top decile of AQP4 expression and high-atrophy re-gions as parcels in the top decile of atrophy. For each high-atrophy parcel, we compute the distance to its nearest AQP4 hotspot using two metrics: (i) Euclidean distance in anatomical space and (ii) shortest path distance along the structural connectome (SC), motivated by empirical findings and computational models suggesting that neurodegenerative vulnerability propagates along white-matter connections [22, 29, 57, 96, 109] (Fig. 4a). Shortest path distances are computed on a group-averaged structural connectome derived from the Human Connectome Project (HCP) S1200 release (*N* = 1065; see *Methods*) [121]. We then average distances across high-atrophy parcels to obtain a mean Euclidean and a mean shortest path distance for each disease, and assess significance using Moran spectral randomization [125] (Fig. 4b).

**Figure 4.**
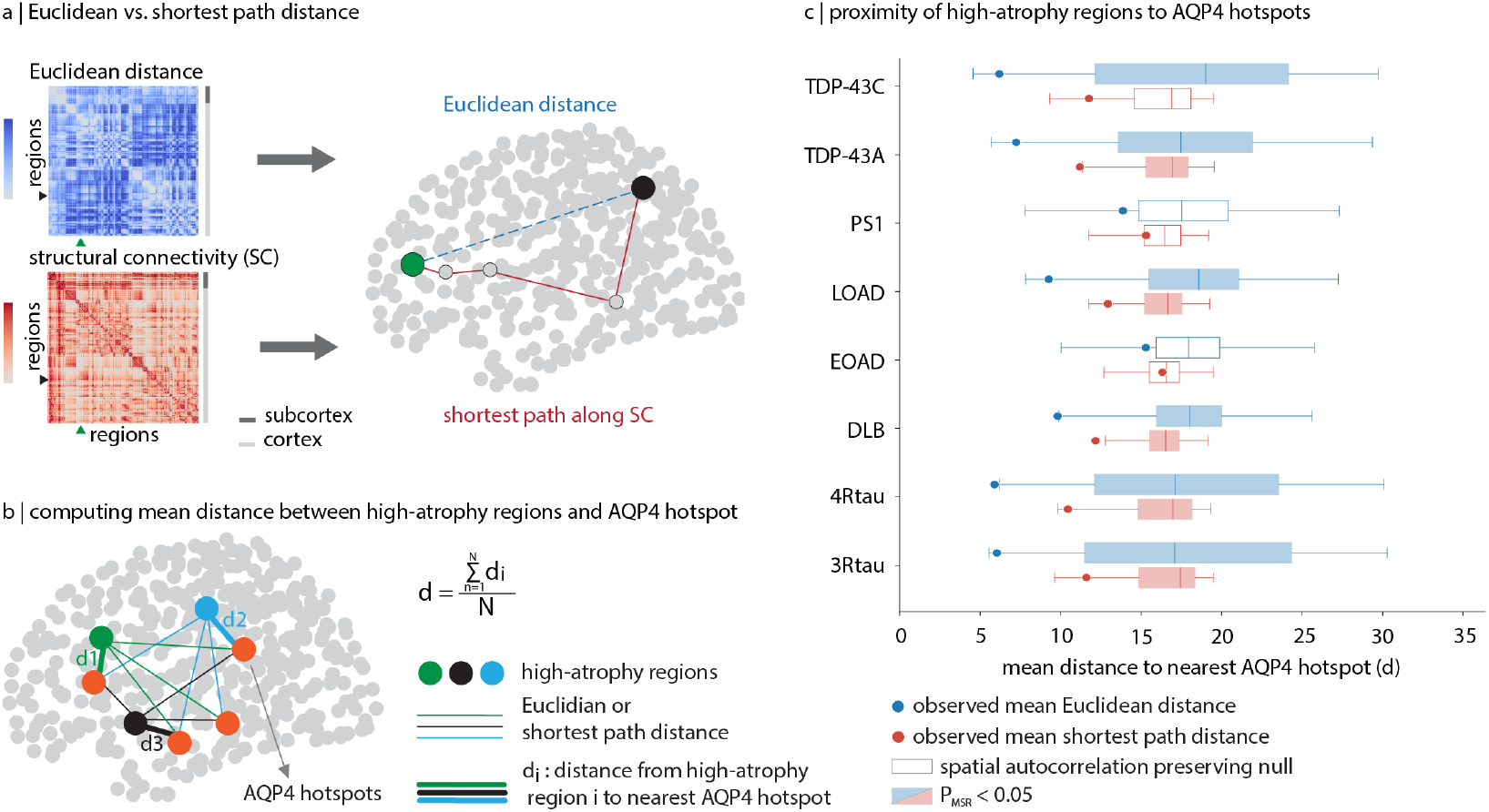
Proximity of high-atrophy regions to AQP4 hotspots in anatomical and network space. (a) Schematic contrasting Euclidean distance in anatomical space with shortest path distance along the SC. (b) Schematic of the proximity analysis. For each disease, high-atrophy regions are parcels in the top decile of disease-related atrophy, and AQP4 hotspots are parcels in the top decile of AQP4 expression. For each high-atrophy parcel, we compute the distance to its nearest AQP4 hotspot using both metrics and average across high-atrophy parcels to obtain a mean distance (*d*) per disease. The SC is computed from the HCP S1200 release (*N* = 1065; see *Methods*) [121]. (c) Observed mean distances (points; blue: Euclidean, red: SC shortest path) are shown relative to spatial autocorrelation–preserving null distributions generated with Moran spectral randomization (*N*_MSR_ = 10 000). Filled boxplots indicate significant proximity to AQP4 hotspots (FDR-corrected, *p*_MSR_ < 0.05). Disease abbreviations match Fig. 3.

For all diseases except PS1 and EOAD, the mean Euclidean distance from high-atrophy parcels to the nearest AQP4 hotspot is significantly reduced relative to the null distribution, indicating that atrophy tends to occur in the spatial neighborhood of AQP4enriched territories. Spatial proximity (Euclidean distance) is significant for TDP-43C (*d* = 6.19; FDR-corrected, *p*_MSR_ = 11.5 *×* 10^−3^), TDP-43A (*d* = 7.24; FDR-corrected, *p*_MSR_ = 7.68 *×* 10^−3^), LOAD (*d* = 9.26; FDR-corrected, *p*_MSR_ = 7.68 *×* 10^−3^), DLB (*d* = 9.81; FDR-corrected, *p*_MSR_ = 2.93 × 10^−3^), 4Rtau (*d* = 5.89; FDR-corrected, *p*_MSR_ = 1.59 ×10^−3^), and 3Rtau (*d* = 6.04; FDR-corrected, *p*_MSR_ = 1.59 × 10^−3^), but not for PS1 (*d* = 13.86; FDR-corrected, *p*_MSR_ = 0.182) or EOAD (*d* = 15.29; FDR-corrected, *p*_MSR_ = 0.182) (Fig. 4c). We find similar results in network space. Shortest path proximity is significant for TDP-43A (*d* = 11.19; FDR-corrected, *p*_MSR_ = 0.036), LOAD (*d* = 12.94; FDR-corrected, *p*_MSR_ = 0.048), DLB (*d* = 12.17; FDR-corrected, *p*_MSR_ = 0.019), 4Rtau (*d* = 10.45; FDR-corrected, *p*_MSR_ = 0.019), 3Rtau (*d* = 11.59; FDR-corrected, *p*_MSR_ = 0.048), but not for TDP-43C (*d* = 11.75; FDR-corrected, *p*_MSR_ = 0.071), PS1 (*d* = 15.31; FDR-corrected, *p*_MSR_ = 0.311), and EOAD (*d* = 16.32; FDR-corrected, *p*_MSR_ = 0.433) (Fig. 4c). Together, these results show that AQP4 hotspots and high-atrophy regions are proximal in both anatomical space and along structural network paths, consistent with the notion that neurodegenerative disease spreads trans-synaptically [22, 29, 57, 96, 109].

### Inflammation modulates the link between AQP4 and neurodegeneration

The relationship between the glymphatic system and neurodegeneration is thought to be intertwined with neuroinflammation. Experimental work suggests that neuroinflammatory processes can perturb AQP4 perivascular polarization, slow glymphatic clearance, and promote protein aggregation [15, 47]. These findings raise the possibility that the brain-wide inflammatory milieu helps shape where atrophy is expressed, and that inflammation-related topography may provide information about regional vulnerability that complements AQP4-linked glymphatic organization. Accordingly, we test whether normative inflammation markers explain variance in disease-related atrophy beyond AQP4 expression, thereby strengthening the alignment between AQP4 topography and neurodegenerative patterns.

To capture spatial variation in neuroinflammatory tone, we use normative *in vivo* positron emis-sion tomography (PET) maps of translocator protein (TSPO) [70], cyclooxygenase-1 (COX-1) [60], and cyclooxygenase-2 (COX-2) [130] in healthy adults. TSPO is an outer mitochondrial membrane protein whose PET signal is widely used as an index of neuroimmune activation, often attributed to activated glial responses, but it is not strictly cell-specific and can reflect contributions from multiple cellular compartments depending on context [37, 64]. COX-1 and COX-2 are key enzymes in prostaglandin synthesis; COX-1 is more constitutive and is linked to microglial (innate immune) signaling, whereas COX-2 is an inducible prostaglandin pathway that is prominent in neurons and upregulated by stress and pathology [60, 130]. Together, these tracers provide complementary molecular indices of inflammation-related signaling, allowing us to test whether inflammatory tone modulates regional vulnerability linked to AQP4.

To test whether inflammatory markers improve the alignment between AQP4 expression and atro-phy, we first fit a simple linear model in which atrophy is predicted from AQP4 expression alone, and quantify model fit using adjusted 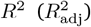. We then extend the model by adding either TSPO, COX1, or COX-2 as an additional predictor and compute the change in model fit, 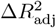. Improvements in model performance are tested against a null model in which the values of the additional predictor are randomized while preserving spatial autocorrelation using Moran spectral randomization [125]. Any increases are therefore not due to the addition of a predictor or its spatial smoothness. Significant improvements are highlighted in Fig. 5a–c.

**Figure 5.**
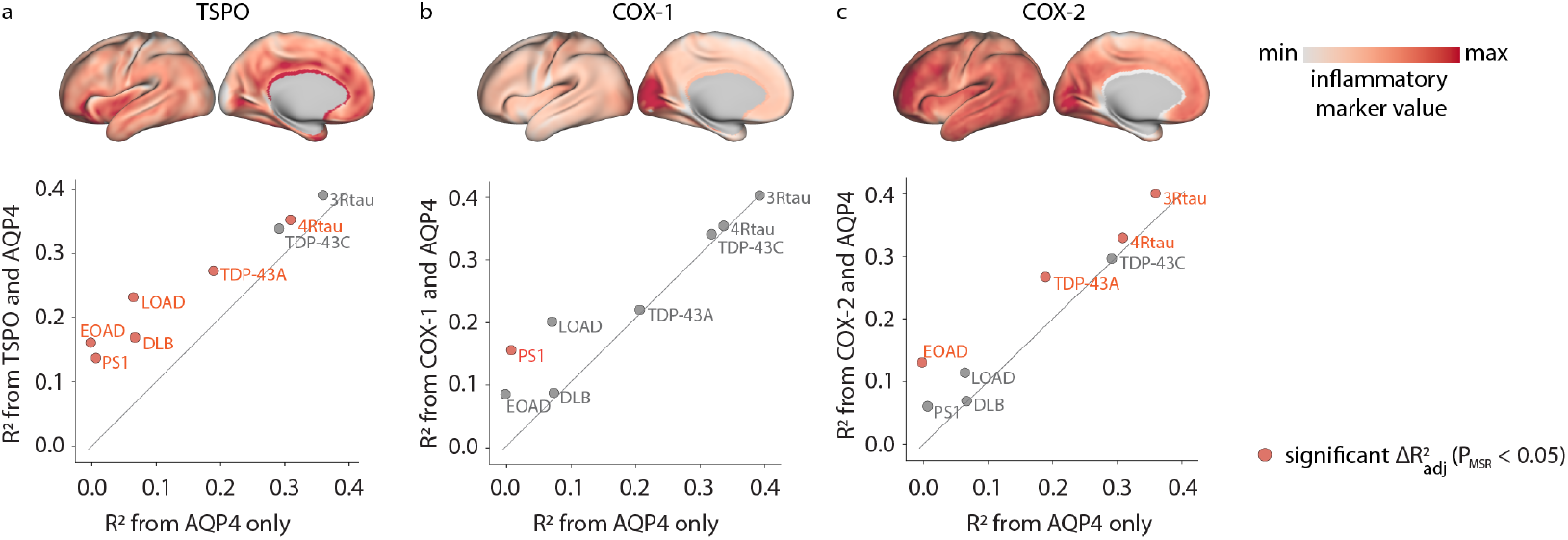
Inflammatory markers augment AQP4–atrophy associations. (a–c) Top: normative PET-derived maps in healthy adults for neuroinflammatory markers including TSPO [70], COX-1 [60], and COX-2 [130] (from left to right). Bottom: for each disease, we quantify model fit for an AQP4-only model (x-axis) and for a model that additionally includes TSPO (a), COX-1 (b), or COX-2 (c) (y-axis), using adjusted 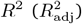. Each point is a disease; the identity line is shown in grey. Points above the line indicate improved fit after adding the marker. Orange points indicate significant improvements in fit relative to spatial autocorrelation–preserving null models generated with Moran spectral randomization (*p*_MSR_ < 0.05) [125]. Disease abbreviations match Fig. 3.

Figure 5 summarizes results across diseases. Across markers, most points fall above the identity line, indicating that adding an inflammatory marker generally improves the prediction of atrophy beyond AQP4 alone. The clearest and most consistent improvements are seen for TSPO and COX-2. In particular, TSPO produces marked gains for the Alzheimer’s spectrum and Lewy body disease (EOAD, LOAD, PS1, DLB), where the AQP4-only model explains relatively little variance but model fit increases substantially once TSPO is included (Fig. 5a). By contrast, COX-2 preferentially boosts fit for proteinopathies with stronger baseline coupling, most notably tauopathies and TDP-43A (Fig. 5c). In comparison, COX-1 yields smaller and less systematic changes in model fit (Fig. 5b). Together, these findings suggest that inflammatory markers provide information about regional vulnerability that is not captured by AQP4 expression alone, strengthening the alignment between glymphatic-related topography and disease atrophy patterns.

### AQP4 channels and susceptibility to edema

Cerebral edema is a common complication of glioma. It reflects excess fluid accumulation in brain tissue surrounding the tumor, driven in part by abnormal tumor vasculature and blood–brain barrier disruption [87]. Because AQP4 is the dominant water channel in astrocytes and is enriched at perivascular endfeet, it sits at a key interface for water exchange between blood vessels and brain tissue, and therefore can influence edema [129]. In glioma, AQP4 is frequently upregulated and redistributed away from its typical perivascular location, and AQP4 levels in peritumoral tissue scale with the degree of edema [17, 83]. More broadly, growing evidence links glioma to disruption of glymphatic fluid transport and neuroinflammation, suggesting that regional differences in AQP4 expression help shape where peritumoral edema is likely to accumulate [17, 129]. Motivated by these observations, we ask whether AQP4 expression aligns with the distribution of peritumoral edema.

To characterize the spatial distribution of peritumoral edema, we use expert delineations of peritumoral edema from two glioma MRI cohorts (*N* = 501; [16] and *N* = 630; [8]). Individual edema masks are aggregated into a voxel-wise edema frequency map, which we then parcellate to the Schaefer–Melbourne atlas [107, 117] (see *Methods*). Edema frequency is not uniformly distributed across cortex and subcortex (Fig. 6a,b). In cortex, edema frequency is higher in ventral insular and peri-Sylvian regions and sensorimotor cortex, with additional involvement in temporal default-network territories, whereas visual cortex shows consistently low involvement (Fig. 6a). Subcortical edema frequency is highest in basal ganglia territories, peaking in putamen and ventral pallidum and remaining elevated in pallidum, while it is comparatively lower across thalamic and other striatal subdivisions, including nucleus accumbens (Fig. 6b).

**Figure 6.**
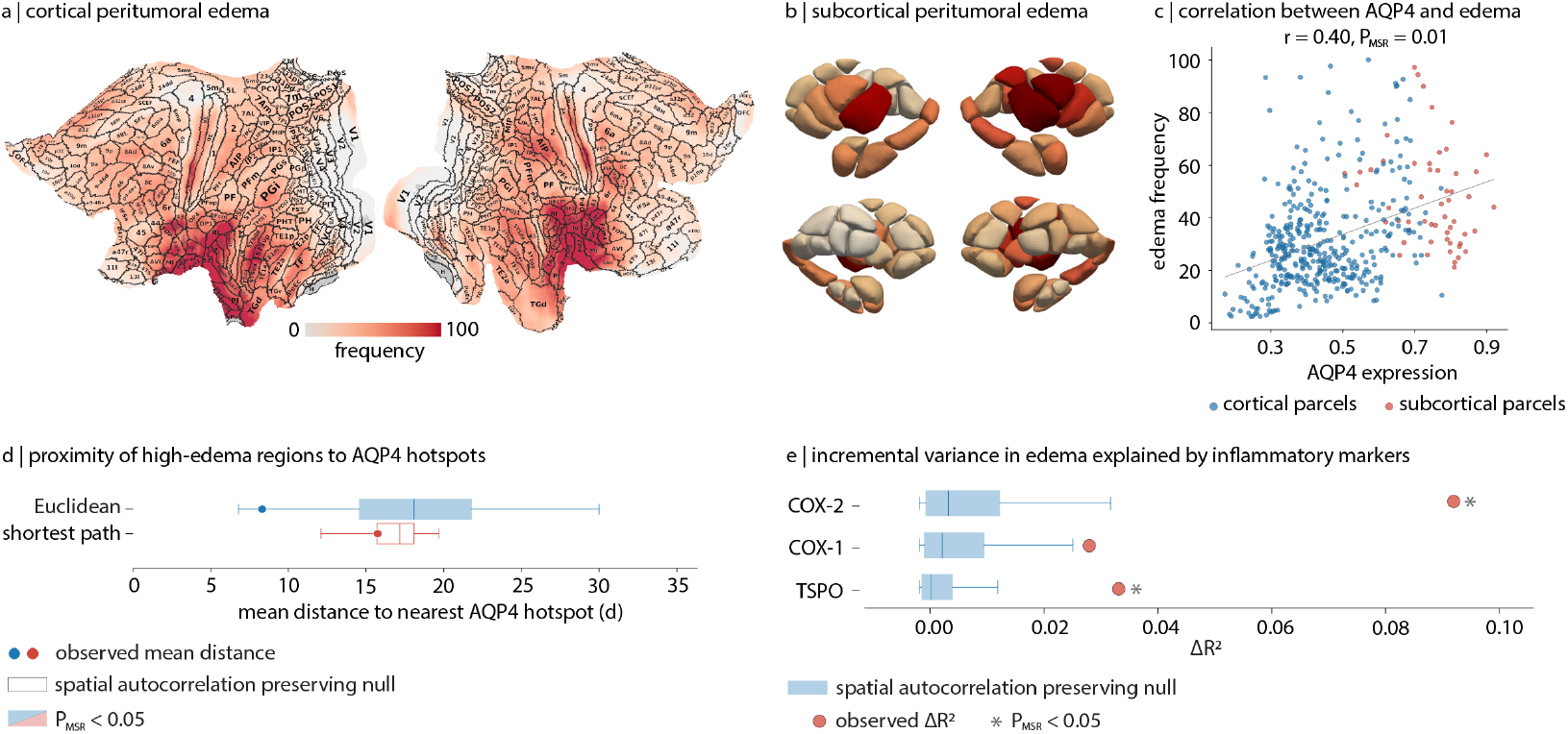
Peritumoral edema aligns with AQP4 expression. (a,b) Normative maps of peritumoral edema frequency estimated from expert edema delineations in two pre-operative glioma MRI cohorts (*N* = 501 [16] and *N* = 630 [8]; see *Methods*). Individual edema masks are aggregated into a voxel-wise edema frequency map and parcellated to the Schaefer–Melbourne atlas [107, 117]. (a) Cortical edema frequency is shown on 2D flat fsLR surfaces with Glasser borders and areal abbreviations overlaid [33] (Table. S2). (b) Subcortical edema frequency is shown in the Melbourne Subcortex S4 Atlas [117] (Table. S1; Fig. S4). (c) Pearson’s correlation between AQP4 expression and edema frequency (*r* = 0.40, *p*_MSR_ = 0.01). (d) Proximity of high-edema parcels (top decile of edema frequency) to AQP4 hotspots (top decile of AQP4 expression), quantified as mean Euclidean distance and mean shortest path distance along the SC. Shortest path distances are computed on a group-averaged connectome derived from the HCP S1200 release (*N* = 1065; see *Methods*) [121]. Observed mean distances (points) are shown relative to spatial autocorrelation–preserving null distributions (boxplots); filled boxplots indicate *p*_MSR_ < 0.05. (e) Incremental variance explained 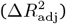 when adding TSPO [70], COX-1 [60], and COX-2 [130] to an AQP4-only model predicting edema frequency. Points show observed 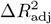 values, and boxplots show spatial autocorrelation–preserving null distributions; asterisks denote *p*_MSR_ < 0.05.

We first ask whether edema frequency co-localizes with AQP4 expression. We find that AQP4 expression is positively correlated with edema frequency (Fig. 6c; *r* = 0.40, *p*_MSR_ = 0.01), indicating that regions with higher AQP4 expression tend to show edema more frequently in these tumor cohorts. Moreover, high-edema parcels show reduced Euclidean distance to AQP4 hotspots relative to spatial autocorrelation–preserving null expectations (Fig. 6d), indicating that edema concentrates in the spatial neighborhood of AQP4-enriched territories.

We then test whether normative PET markers of neuroinflammation (TSPO [70], COX-1 [60], and COX-2 [130]) explain additional variance in edema beyond AQP4. Using the same hierarchical regression framework as above, adding each marker improves model fit relative to AQP4 alone, with the largest increase for COX-2 (Fig. 6e; COX-2: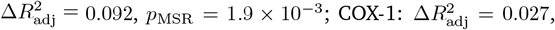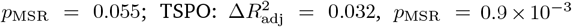. Together, these results suggest that peritumoral edema co-localizes with AQP4 expression and is additionally shaped by the regional inflammatory tone.

### Extending to other aquaporins

Aquaporin-mediated water (and solute) transport in the brain is distributed across multiple isoforms. Among these, AQP1, AQP4, and AQP9 have the most consistent evidence for their roles in brain physiology and disease [4, 68, 114, 132]. AQP1 is highly expressed in the choroid plexus epithelium and has been implicated in facilitating transepithelial water permeability and CSF secretion [88]. In contrast, AQP4 is the predominant aquaporin in the brain parenchyma, enriched in astrocytes and polarized to perivascular endfeet and the glia limitans, positioning it at key CSF–ISF interfaces implicated in perivascular exchange and clearance [50, 84]. AQP9, by comparison, is an aquaglyceroporin expressed in subsets of astrocytes and other neural cell types (including periventricular populations), with permeability to glycerol and other small solutes, linking it more directly to metabolic substrate transport than to purely water-selective flux [5, 6]. Motivated by these distinct physiological roles— and the central role of AQP4 in waste clearance and water homeostasis—we apply the same AHBA-based mapping and whole-brain association framework to AQP1 and AQP9 as within-family comparators alongside AQP4.

First, we ask how AQP4 co-varies with AQP1 and AQP9 across parcels. We find that AQP1 and AQP4 show very similar spatial patterns (*r* = 0.79, *p*_MSR_ = 1.0 × 10^−4^), whereas AQP9 shows an inverse spatial alignment with both AQP1 (*r* = −0.48, *p*_MSR_ = 0.016) and AQP4 (*r* = −0.47, *p*_MSR_ = 2.4 × 10^−3^) (Fig. 7a). We further investigate the spatial association between each aquaporin expression and the phenotypes examined in this study, including vascular measures, dementia atrophy patterns, and peritumoral edema. Across phenotypes, AQP1 shows a similar profile to AQP4; both show negative correlations with vascular measures and positive correla-tions with neurodegenerative atrophy patterns and with peritumoral edema. In contrast, AQP9 shows opposite-sign correlations (Fig. 7b).

**Figure 7.**
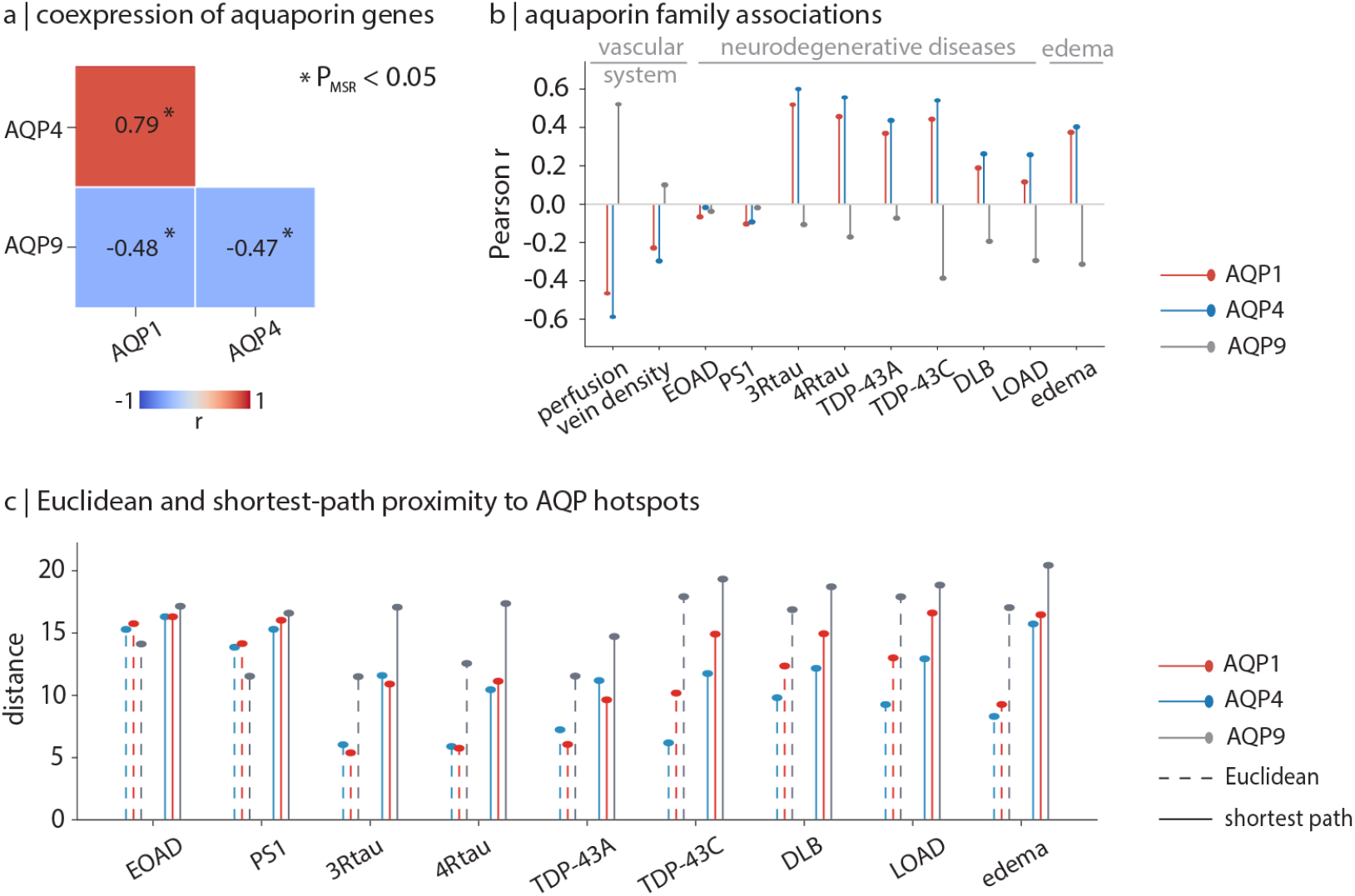
Extending to other aquaporins. (a) Pearson correlations among AHBA-derived expression maps of AQP1, AQP4, and AQP9 [45]. (b) Correlations between each aquaporin expression map and the phenotypes studied here: vascular measures (blood perfusion PC1 [23]; VENAT vein density [48]), neurodegenerative atrophy patterns (EOAD, PS1, 3Rtau, 4Rtau, TDP-43A, TDP-43C, DLB, LOAD) [42], and peritumoral edema frequency [8, 16]. (c) Mean distance from disease vulnerable parcels (top decile of atrophy or edema) to the nearest aquaporin hotspot (top decile of expression), computed using Euclidean distance (dashed) or SC shortest path distance (solid). See Fig. 4a,b for a schematic of the proximity analysis. Gene maps are processed with the abagen workflow [76] and parcellated to the Schaefer–Melbourne atlas (see *Methods*) [107, 117].

We next ask whether vulnerable territories are preferentially located near aquaporin-enriched regions using the same hotspot proximity framework as for AQP4. For each gene, hotspots are defined as the top decile of expression, and we quantify the mean distance from high-atrophy (or high-edema) parcels to the nearest hotspot using both Euclidean distance and SC shortest path distance. Figure 7c shows that vulnerable territories are generally closer to AQP4 and AQP1 hotspots than to AQP9 hotspots. Together, these results suggest that the spatial coupling observed for AQP4 is not shared uniformly across the aquaporin family.

## DISCUSSION

In this study, we provide a whole-brain account of glymphatic organization using AQP4 expression as a molecular anchor. We first map AQP4 expression across cortex and subcortex using AHBA microarray data. We find the highest expression in periventricular and ventral subcortical regions, and in limbic and transmodal cortices. We next relate this spatial axis to vascular organization, neurodegenerative vulnerability, neuroinflammation, and edema. First, AQP4 expression is anticorrelated with normative perfusion and with a vein density atlas, suggesting that glymphatic-related territories are not sim-ply a reflection of vascular supply or large-vessel anatomy. Second, AQP4 expression co-localizes with atrophy patterns across several pathologically confirmed dementias, most strongly in tau and TDP-43 proteinopathies, and these associations often strengthen when inflammatory markers are incorporated. Third, peritumoral edema frequency aligns with AQP4 expression and is further explained by inflammatory markers. Finally, comparisons within the aquaporin family suggest that these effects are not shared uniformly across isoforms. AQP4 (and, to a lesser extent, AQP1) shows consistent coupling to the phenotypes examined here, whereas AQP9 follows a distinct spatial profile. Together, our findings place AQP4 at the intersection of perivascular fluid exchange, vascular physiology, inflammation, and vulnerability to neurodegeneration and edema.

AQP4 is enriched in astrocytes and shows strong subcellular polarization to perivascular endfeet and the glia limitans, placing it at major CSF–ISF interfaces implicated in perivascular exchange [38]. In this context, the high AQP4 expression we observe in periventricular and ventral subcortical territories—including basal ganglia and amygdalar regions—is consistent with the broader view that CSF–ISF exchange involves both perivascular routes and interfaces adjacent to CSF reservoirs, including ventricular surfaces [63, 106]. In cortex, higher expression in limbic and transmodal areas and lower expression in primary sensory regions suggests that the glymphatic system is organized along gradients of cortical organization. Consistent with this idea, the diffusion tensor imaging–analysis along the perivascular space (DTI-ALPS) index shows a clear large-scale topography, with developmental changes following a posterior–anterior axis and the strongest age-related increases moving toward anterior regions [69]. More broadly, emerging work shows that vascular and perivascular cell properties, including astrocytic and endothelial specializations, vary systematically across brain regions [94]. This supports the idea that perivascular astrocyte specializations—and, by extension, aspects of regional fluid-handling capacity—may be distributed in a principled, non-uniform manner rather than being spatially homogeneous.

Glymphatic transport is anatomically embedded in the cerebrovascular tree. Perivascular spaces surround penetrating vessels, and CSF–ISF exchange along these pathways is modulated by multiple physiological drivers, including arterial pulsatility, respiration-linked pressure changes, and slow vasomotion [44, 51, 81, 116, 122]. Despite this coupling, we find that AQP4 expression is anticorrelated with cerebral perfusion and with a vein density atlas. A parsimonious interpretation is that perfusion and vein density mainly reflect blood supply and large-vessel anatomy, whereas perivascular exchange depends more on perivascular spaces around penetrating small vessels and their local proper-ties, including their geometry and the polarization of astrocytic endfeet that regulate water transport [52, 98, 106, 126, 136]. In this view, vascular supply and glymphatic-related tissue organization impose partly distinct spatial constraints on brain physiology.

A major motivation for studying glymphatic organization is the long-standing hypothesis that regional neurodegenerative vulnerability is shaped, in part, by how efficiently interstitial solutes can be cleared along glymphatic outflow routes. One possibility is that brain regions with high AQP4 expression are a bottleneck for glymphatic outflow, making them natural sites where solutes and aggregation-prone proteins can accumulate over time and thereby increasing susceptibility to atrophy. Another possibility is that impaired perivascular clearance contributes to the accumulation and spread of pathogenic proteins in neurodegeneration [50, 85, 97]. This idea is supported by experimental evidence showing that reduced glymphatic exchange slows amyloid-*β* clearance and can exacerbate amyloid burden, and that disrupting AQP4 function or perivascular AQP4 localization impairs exchange and promotes amyloid pathology [80, 104, 110]. Parallel work supports a role for glymphatic pathways in tau handling, with impaired glymphatic function reducing tau clearance and facilitating tau accumulation in disease models [43, 49, 53, 71]. Moreover, human studies further suggest that glymphatic dysfunction is graded and heterogeneous across individuals and brain regions, and that reactive astrogliosis and loss of AQP4 polarization may contribute to impaired perivascular exchange in aging and disease [65, 73, 106, 133]. Our findings complement these studies by showing that, even without direct measures of perivascular polarization or glymphatic flux, regional AQP4 expression systematically aligns with where neurodegeneration occurs across a range of pathologically confirmed dementias, most strongly for tau and TDP-43 proteinopathies. One interpretation is that AQP4-rich territories coincide with key waste clearance bottlenecks where proteins accumulate most, while a second, non-exclusive interpretation is that these territories place greater demand on intact AQP4-supported waste clearance, such that depolarization or dysfunction of AQP4 disproportionately compromises clearance and increases long-term vulnerability.

Beyond map-to-map correlations, we test whether high-atrophy territories tend to lie near AQP4-enriched regions. We find that, for most diseases, high-atrophy regions are close to AQP4 hotspots, both in Euclidean space and along shortest paths along the structural connectome. This suggests that neurodegeneration tends to cluster around AQP4-rich regions, not only in anatomical space but also in network space. More broadly, this pattern fits network-based models of neurodegeneration, where local susceptibility interacts with connectome topology and network architecture shapes the spatial patterning of disease effects [22, 29, 57, 96, 109]. In this view, AQP4-rich regions may mark territories where impaired clearance and local proteostatic stress are most likely to initiate or amplify damage, while connections provide routes through which vulnerability can propagate to nearby regions in network space. However, proximity does not imply causality. AQP4 may also covary with other macroscale axes such as cytoarchitecture, myelination, or cortical hierarchy [10, 14, 35]. Future work should test whether AQP4 explains additional variance in disease topography beyond these established gradients.

Additionally, our study finds that neuroinflammatory topography refines the association between AQP4 and neurodegeneration. Across diseases, adding normative PET markers of inflammatory tone (TSPO, COX-1, COX-2) typically improves the explanatory power of AQP4 for atrophy, with the most consistent gains for TSPO and COX-2. This fits a view in which perivascular transport, vascular permeability, and inflammatory signaling are linked. Inflammatory activation can shift astrocyte state, disrupt endfoot organization, and reshape perivascular spaces, which can impair solute clearance. Conversely, reduced clearance can prolong exposure to extracellular debris and misfolded proteins, which can sustain neuroimmune activation [15, 30, 47, 52]. The marker-specific patterns we observe may also reflect differences in what these tracers capture. TSPO PET is widely used to index neuroimmune activation, but it is not cell-specific and can reflect multiple cellular sources depending on context [37, 64, 86]. COX-1 is often treated as a more constitutive prostaglandin pathway with strong myeloid/microglial contributions, whereas COX-2 is prominent in neurons and is inducible by activity and stress [32, 58, 59].

Edema provides a complementary phenotype that is directly tied to brain water balance. In glioma, peritumoral edema is largely vasogenic, reflecting extracellular fluid accumulation driven by blood– brain barrier disruption [87]. In this view, AQP4 is especially relevant because it is the dominant water channel and is enriched at perivascular endfeet, where it regulates water flux at vessel–brain interfaces [30, 62, 90]. Our results show that peritumoral edema frequency is higher in regions with higher AQP4 expression, suggesting that tumor-associated fluid accumulation follows the same topography that organizes AQP4. This aligns with the idea that baseline regional differences in perivascular water-handling machinery may bias where vasogenic edema accumulates once vascular permeability is increased by the tumor, consistent with evidence that AQP4 regulates perivascular water flux and is altered in peritumoral tissue in association with edema [17, 83]. We also find that inflammatory markers explain additional variance in edema beyond AQP4 alone. The largest incremental gain is for COX-2, consistent with evidence that COX-2–dependent prostaglandin signaling can increase vascular permeability and contribute to tumor-associated edema [7]. Together, these findings suggest that tumor-driven vascular leakage, together with AQP4 topography and regional inflammatory tone, shape the spatial pattern of peritumoral edema.

We finally investigate whether these vulnerability associations are specific to AQP4 by repeating the analyses for related aquaporins, AQP1 and AQP9. AQP1 is enriched in the choroid plexus and is commonly linked to CSF secretion [88, 118]. We find that AQP1 largely tracks AQP4 across phenotypes, whereas AQP9, an aquaglyceroporin with closer ties to solute and metabolic substrate transport, shows an opposite-sign association profile across phenotypes. These results suggest that the observed coupling to vulnerability is more specific to water transport and CSF–ISF exchange pathways than to aquaporin expression in general [3, 6, 50, 84, 91, 98].

Measuring glymphatic function in humans remains challenging, and existing *in vivo* methods trade off invasiveness, spatial specificity, and scalability. A common non-invasive option is the DTI-ALPS index, which estimates water diffusion along perivascular spaces from a few periventricular ROIs and is typically reported as a single subject-level measure, not a whole-brain map [115]. More direct *in vivo* evidence comes from intrathecal tracer MRI (often gadobutrol), which measures regional tracer enrichment and delayed clearance; however, this technique is invasive and requires repeated scans over long time windows. Here we use AQP4 expression as a brain-wide molecular axis related to perivascular water exchange, following a broader strategy in which gene transcription is used to map distributed molecular systems at the macroscale (e.g., neuropeptide receptor families) [18, 95]. This atlas-style approach is also consistent with observations in NMOSD, where brain lesions preferentially occur in regions with high AQP4 expression, supporting the idea that AQP4 topography is informative for where AQP4-mediated pathology manifests in the brain [1]. Encouragingly, we find a positive spatial correlation between AQP4 expression and gadobutrol tracer MRI measures from Ringstad and colleagues [101].

While our study offers promising insights, several methodological limitations should be noted. First, we use bulk microarray gene expression data from the Allen Human Brain Atlas (AHBA), which, despite being the most spatially comprehensive dataset currently available, is limited in resolution, sample size, and may not perfectly correspond to protein density [39, 72]. Although we validate the AQP4 spatial pattern against AHBA RNA-seq and the Human Protein Atlas, the extent to which it captures inter-individual variability remains uncertain. Second, glymphatic function depends on perivascular AQP4 polarization and channel organization (including isoform balance and formation of orthogonal arrays), which are not measured here [84]. Third, our analyses use maps from different cohorts and modalities (ASL, QSM, PET, VBM, and tumor MRI), so we cannot infer within-subject associations among perfusion, inflammation, AQP4, and vulnerability. Fourth, the structural connectome used in this study is reconstructed from diffusion-weighted imaging, which is prone to false positives and false negatives [25, 74]. Although we minimize this limitation by employing modern multi-shell diffusion data and state-of-the-art tractography, the inherent challenges of tractography remain. Lastly, disease maps are cross-sectional summaries of atrophy and do not capture longitudinal progression, which limits inference about propagation mechanisms.

Looking ahead, we hope to get all these measures (glymphatic function, perfusion, inflammation, and vulnerability) in the same individuals. In parallel, postmortem histology and single-cell measurements could clarify how regional AQP4 transcription relates to protein abundance, isoform composition, and perivascular polarization across the lifespan and in disease. Furthermore, longitudinal neurodegenerative cohorts will be valuable to test whether alignment to AQP4 predicts future spread, whether inflammatory markers identify periods of accelerated vulnerability, and whether interventions that restore AQP4 polarization or modulate inflammation preferentially benefit AQP4-enriched territories. We make our code and data openly available to facilitate further research and validation of our results.

## METHODS

All code and data used in this study are available at https://github.com/netneurolab/taheri_aquaporin-four.

### Gene expression data

The expression of aquaporin genes was obtained from 6 postmortem brains (1 female, ages 24–57 years, mean ± SD = 42.5 ± 13.4, three of Caucasian ethnicity, two African American, one Hispanic, [45]) using the abagen toolbox (v0.1.4; https://github.com/rmarkello/abagen; [76]). Detailed information about data acquisition can be found in [45]. Preprocessing of gene expression data followed the procedures described in [2]. Microarray probes for all individuals were fetched from https://human.brain-map.org/ and re-annotated to match the gene symbol ID and name from the latest version of NCBI [24], available at https://ftp.ncbi.nih.gov/gene/DATA/GENE_INFO/Mammalia/. Probes not matched to a valid ID were discarded. The re-maining probes underwent intensity-based filtering, where probes with intensity less than background noise in less than 20% of samples across individuals were excluded. For genes represented by multiple probes, the probe with the highest average Pearson correlation to the RNA-seq data from two individuals in the dataset was selected. This ensured biological plausibility of the microarray measurements wherever possible [82]. Tissue sampling locations from the original T1w scans of individuals were non-linearly registered to MNI152 (1mm) space as in https://github.com/chrisgorgo/alleninf. Probe samples were then assigned to brain regions in a standard atlas consisting of the Schaefer 400 cortical [107] and the Melbourne Subcortical S4 [117], thresholded at 0.5 probability. Samples were then assigned to a brain region with MNI coordinates within a 2 mm vicinity. To reduce the potential for misassignment, sample-to-region matching was constrained by hemisphere and gross structural divisions (i.e., cortex, subcortex/brainstem, and cerebellum), such that, e.g., a sample in the left cortex could only be assigned to an atlas voxel in the ipsilateral side. If a brain region was not assigned a sample based on that procedure, voxels in that region were mapped to the nearest tissue sample from the same individual to produce a dense, interpolated expression map. The mean of these expression values was taken across all voxels in the region, weighted by the distance between each voxel and the sample mapped to it, in order to obtain an estimate of the parcellated expression values for the missing region. Inter-subject variation was addressed by normalizing sample expression values across genes using a robust sigmoid function as in [31]. Normalized expression values were then rescaled from 0 to 1 by min-max normalization. Gene expression values were subsequently normalized across regions using an identical procedure. All available tissue samples were used for normalization, whether or not they were assigned to a brain region. Tissue samples not matched to a brain region were discarded after normalization. Samples assigned to the same brain region were averaged separately for each individual, which resulted in six expression matrices, one for each individual, with rows corresponding to brain regions and columns corresponding to genes. From these initial expression matrices, we retained genes with a differential stability value greater than 0.1 [14]. All matrices were finally mean-averaged across individuals, resulting in a single matrix representative of the expression level of a particular gene in a given region.

For visualization only, a dense AQP4 expression was generated in MNI152-NonLinear2009cAsym space using nearest-neighbor interpolation with the abagen workflow. This volumetric map was then projected to fsLR surface space for cortical display. All statistical analyses were performed on regionwise expression values as described above.

### Neurodegenerative disease atrophy data

Data for eight neurodegenerative diseases were taken from the voxel-based morphometry (VBM) maps released by Harper et al. [42] (available via NeuroVault:https://neurovault.org/collections/ADHMHOPN/). The dataset comprises T1-weighted MRI from 186 participants with a clinical dementia diagnosis and histopathological confirmation of the underlying disease (postmortem or biopsy), as well as 73 healthy controls. Condition-level maps reflect averages within each diagnostic group: Alzheimer’s disease (AD; *N* = 107; early-onset *N* = 68, lateonset *N* = 29, PSEN1 mutation carriers *N* = 10), dementia with Lewy bodies (DLB; *N* = 25), three-repeat tauopathy (3Rtau; *N* = 11), four-repeat tauopathy (4Rtau; *N* = 17), FTLD-TDP type A (TDP-43A; *N* = 12), and FTLD-TDP type C (TDP-43C; *N* = 14). Scans were acquired across multiple sites and vendors (Philips, GE, Siemens) with heterogeneous protocols; field strengths included 1.0T (*N* = 15), 1.5T (*N* = 201), and 3T (*N* = 43). Neuropathological assessment followed the standard criteria used at the time of evaluation at one of four centers (Queen Square Brain Bank, King’s College Hospital, VU Medical Center Amsterdam, and the Institute for Ageing and Health, Newcastle) [42].

Regional tissue loss was quantified using VBM on T1-weighted images and reported as voxelwise *t*-statistics. The resulting atrophy maps were adjusted for age, sex, total intracranial volume, scanner field strength, and acquisition site. The volumetric atrophy maps were parcellated to the Schaefer 400 cortical atlas [107] combined with the Melbourne sub-cortical S4 atlas [117]. Ethics approval for the retrospective study is described in Harper et al. [42].

### Structural connectivity data

The template diffusion-weighted MRI (dMRI) connectome used to compute structural shortest paths was derived from 1065 participants in the Human Connectome Project (HCP) S1200 release. dMRI data were acquired with a spin–echo echoplanar imaging (EPI) sequence (TR = 5520 ms, TE = 89.5 ms, flip angle = 78^°^, 1.25 mm isotropic voxels) using *b*-values of 1000, 2000, and 3000 s/mm^2^. Three diffusion runs with different gradi-ent direction sets were collected (90 directions total). Full acquisition details are reported elsewhere [21, 121]. Preprocessing followed the HCP Minimal Preprocessing Pipelines [34], including intensity normalization, susceptibility distortion correction (topup), eddy-current and head-motion correction (eddy), and gradient nonlinearity correction. We downloaded bedpostX-derived fibre orientation density function (fODF) estimates from the HCP S1200 release and performed probabilistic tractography with `probtrackx2` to estimate interregional structural connectivity (QuNex v0.98.0, `dwi_probtrackx_dense_gpu;` [56]) using Schaefer 400 cortical atlas [107] combined with the Melbourne subcortical S4 atlas [117].

### Shortest path retrieval

Structural connectome was encoded as an undirected weighted graph *G* ≡ {*V, W*} composed of nodes *V* = {*v*_1_, *v*_2_, …, *v*_*n*_} and a matrix of fiber density values *W* = [*w*_*ij*_], valued on the interval [0, 1]. To recover shortest paths, we first define a topological distance measure. The weighted adjacency matrix was transformed from connection weights to connection lengths using the transform *L* = 1*/W*, such that connections with greater weights are mapped to shorter lengths [36]. Note that other transformations are also possible, including *L* = − log(*W*). Weighted shortest paths were recovered using the Floyd–Warshall algorithm ([28, 105, 127]; *BrainConn* Python Toolbox). Note that in many types of networks there may exist multiple shortest paths between two nodes (edge-disjoint or not); in our network, this was not the case, as we computed weighted shortest paths yielding unique paths between all source–target pairs.

### Spatial autocorrelation–preserving null models

Many of the maps used in this study (AQP4 expression, perfusion, venous density, inflammation biomarkers, and disease atrophy) exhibit strong spatial autocorrelation. Standard parametric tests, therefore, overestimate the effective degrees of freedom [78, 123]. To obtain more appropriate significance estimates, we assessed all spatial associations against null models that preserve the empirical spatial autocorrelation using Moran spectral randomisation (MSR), as implemented in the BrainSpace toolbox (https://brainspace.readthedocs.io/en/latest/) [124].

For each analysis, we generated 10,000 surrogate maps with matched spatial smoothness. MSR first quantifies spatial autocorrelation in the empirical map using Moran’s *I*, defined with a spatial weight matrix based on the inverse Euclidean distance between parcel centroids in MNI space. The corresponding spatial eigenvectors (Moran eigenvector maps) provide an orthogonal basis that captures the spatial dependence structure of the data. Surrogate maps are then obtained by sampling random coefficients in this eigenvector basis and recombining them such that the resulting maps retain the same overall spatial autocorrelation as the empirical map, while otherwise being random [78, 124].

For pairwise correlations (e.g. AQP4 vs perfusion, venous density, or atrophy), we held one map fixed and applied MSR to the other map to obtain a null distribution of correlation coefficients. For the proximity analyses, we generated MSR surrogates of the disease or edema map, redefined high atrophy or high edema for each surrogate, and recomputed Euclidean and shortest path distances. Unless otherwise noted, *p*_MSR_ values are one-sided in the direction of the observed effect.

### Blood perfusion data

Cerebral blood perfusion data was obtained from the Human Connectome Project Lifespan studies (HCP-Development and HCP-Aging), comprising 1305 healthy participants spanning childhood to late adulthood [13, 41, 112]. Perfusion is estimated using pseudo-continuous arterial spin labeling (pCASL) magnetic resonance imaging acquired on a 3T Siemens Prisma scanner with a 32-channel head coil. The pCASL sequence used a 2D multiband echo-planar imaging readout with a labeling duration of 1500 ms and five post-labeling delays (200, 700, 1200, 1700, and 2200 ms), yielding 6, 6, 6, 10, and 15 control–label pairs at each delay. Additional parameters included isotropic voxel size of 2.5 mm and TR/TE of 3580/18.7 ms, together with two M_0_ calibration images for quantification in ml/100 g/min.

ASL data were processed with the HCP ASL pipeline as implemented in QuNex [56]. Label, control, and calibration images were co-registered to each subject’s anatomical space, and motion, susceptibility distortion, and gradient nonlinearity effects were corrected in a single resampling operation. We then applied bias-field and banding artifact corrections and performed motion-aware label– control subtraction. Cerebral blood flow and arterial transit time were computed in `oxford_asl` using a multi-compartment Buxton model, with partial-volume effects handled via a spatial variational Bayes procedure. Quantitative perfusion maps were subsequently projected to grayordinates (cortical surface plus subcortical voxels).

To derive a normative perfusion topography shared across participants, we summarized individual perfusion maps using principal component analysis (PCA), following prior work [23]. Briefly, participant-level perfusion maps were z-scored and concatenated into a participant-by-grayordinate data matrix. This participant-wise standardization reduces global offsets in perfusion levels (e.g., differences related to biological sex) and emphasizes shared spatial patterning. PCA was then applied to this matrix, and the first principal component (PC1) was retained as the group-level “perfusion score” map, as it captures the dominant regional pattern of perfusion shared across individuals (PC1 explained 50.71% of the variance; PC2 explained 2.35%). The resulting group perfusion map was then parcellated to the Schaefer 400 cortical [107] and Melbourne subcortical S4 [117] atlases to obtain regional perfusion estimates.

### Vein density data

Normative atlas of cerebral veins was obtained from the Venous Neuroanatomy (VENAT) dataset [48]. VENAT was built from 7 T quantitative susceptibility mapping (QSM) acquired at 0.6 mm isotropic resolution in healthy young adults, with five repeated scans per participant (*N* = 20; mean age 25.1 ± 2.5 years). Veins were segmented in 3D and aligned across participants using a two-step registration procedure. The atlas provides voxel-wise maps of average vessel location, which we treat as a vein density (or vein probability) map, and also includes summary maps of vessel diameter (mean 0.84 ± 0.33 mm) and curvature (mean 0.11 ± 0.05 mm^−1^). In our analyses, we used the average vessel location map as the vein density phenotype. We parcellated it to the Schaefer 400 cortical atlas combined with the Melbourne subcortical S4 atlas [107, 117].

### Peritumoral edema data

Peritumoral edema maps were derived from two large, publicly available pre-operative glioma MRI resources: the UCSF Preoperative Diffuse Glioma MRI (UCSF-PDGM) dataset [16] and the University of Pennsylvania glioblastoma (UPenn-GBM) cohort [8]. UCSF-PDGM includes 501 adults with diffuse gliomas spanning World Health Organization (WHO) grades 2–4 (higher grade indicates more aggressive disease), imaged with a standard-ized pre-operative 3T MRI protocol and accompanied by multi-compartment tumor segmentations. In this dataset, we used the *FLAIR abnormality* compartment—a non-enhancing T2-FLAIR hyperintense region that includes edema and infiltrative tumor—as our edema mask [16]. UPenn-GBM includes 630 patients with de novo glioblastoma and provides automated tumor sub-region annotations that were manually revised. We used the *peritumoral edematous* (ED) label, which likewise corresponds to the hyperintense peritumoral envelope on T2-FLAIR [8]. In both datasets, we treated the T2-FLAIR hyperintense peritumoral compartment (ED / FLAIR abnormality) as a proxy for peritumoral edema.

To obtain a group-level map of edema distribution, individual edema masks were transformed to MNI152 standard space and resampled to a common voxel grid. We then computed a voxel-wise edema frequency map by summing binary masks across participants, such that each voxel value reflects the number of individuals whose edema mask includes that voxel. Finally, we parcellated the resulting frequency map to the Schaefer 400 cortical atlas [107] combined with the Melbourne sub-cortical S4 atlas [117], yielding a regional edema frequency score for each parcel in the 454-region whole-brain parcellation.

### PET markers of neuroinflammation data

PET-derived maps of neuroinflammatory markers were obtained from normative atlases in healthy adults. We downloaded translocator protein (TSPO) and cyclooxygenase-1 (COX-1) maps from the `neuromaps` toolbox (https://github.com/netneurolab/neuromaps), which curates a library of brain annotations and provides standardized access and transformations across common coordinate systems [77]. Specifically, we used a TSPO PET map derived from [70] (radioligand: [11C]PBR28) and a COX-1 PET map derived from [60] (radioligand: [11C]PS13), as distributed in the `neuromaps` reference dataset. Methodological details regarding tracer acquisition, quantification, and preprocessing are reported in the original studies and in the `neuromaps` documentation (https://netneurolab.github.io/neuromaps/listofmaps.html). Furthermore, a normative COX-2 PET map was obtained from [130] (radioligand: [11C]MC1). Volumetric PET images were parcellated to the Schaefer 400 cortical [107] and Melbourne subcortical S4 [117] atlases to obtain regional estimates.

### Intrathecal CSF tracer enrichment data

As an *in vivo* tracer-based assay of the glymphatic system, we used regional brain enrichment measures from a prior MRI study by Ringstad et al. [101]. In that study, reference participants (REF; *N* = 8, evaluated for suspected CSF leak/idiopathic intracranial hypotension) underwent repeated T1-weighted MRI before and after intrathecal administration of the CSF tracer gadobutrol (0.5 ml; mmol/ml; Gadovist, Bayer). MRI data were acquired on a 3T Philips Ingenia scanner using a consistent 3D T1-weighted protocol across time points. Post-contrast T1-weighted scans were repeated during the first hour, approximately every two hours until late afternoon, and again the next morning (approximately 24 hours post-injection). Scans were subsequently grouped into predefined time windows, including a 24 hour interval.

Image processing was performed in FreeSurfer (v6.0) [26] using a longitudinal workflow with within-subject template construction and rigid alignment across sessions [99]. Tracer-related signal was quantified for each FreeSurfer-derived segmented region by computing the median T1 intensity and normalizing it to a reference ROI placed in the posterior superior sagittal sinus, yielding normalized T1 signal units that mitigate global intensity scaling [101]. Regional tracer enrichment was then expressed as percentage change from the pre-contrast baseline [101].

In the present work, we focus on the REF group at 24 hours because this is a standardized delayed time point in which gadobutrol has distributed broadly beyond the early peri-surface phase, providing a stable snapshot of net parenchymal enrichment suitable for spatial comparison with AQP4 expression [101].

### Human Protein Atlas gene expression data

Human Protein Atlas gene expression measures were obtained from RNA sequencing (RNA-seq) of micropunch samples collected at the Human Brain Tissue Bank, Semmelweis University (Hungary), spanning 190 brain regions, areas, and subfields [111, 119]. RNA was extracted with the RNeasy Plus Mini Kit (Qiagen), and mRNA was enriched via ribosomal RNA depletion. RNA quality was assessed using the Experion RNA High-Sens Analysis kit (Bio-Rad), and only samples passing minimum thresholds for RNA Integrity Number (RIN) and the 260/280 absorbance ratio were retained. Libraries were generated with Illumina TruSeq Stranded mRNA reagents and sequenced on an Illumina NovaSeq 6000 system (paired-end, 150 bp). Reads were aligned to GRCh37/hg19 using Ensembl gene models (v92) and quantified with Kallisto (v0.43.1) [111, 134, 135]. Following quality control, expression was summarized as TPM and then further normalized using trimmed mean of Mvalues (TMM) in NOISeq, with batch effects adjusted using `limma` [102]. Normalized TPM (nTPM) values were log-transformed (log_10_) and assigned to Destrieux atlas regions via region-name matching [20].

## Acknowledgments

We thank Filip Milisav, Vincent Bazinet, Yigu Zhou, Moohebat Pourmajidian, and Maria Haddad for their comments and suggestions on the manuscript. BM acknowledges support from the Natural Sciences and Engineering Research Council of Canada (RGPIN-2017-04265), Canadian Institutes of Health Research (PJT-180439), and Canada Research Chairs Program (CRC-2022-00169).

---

**Figure S1.**
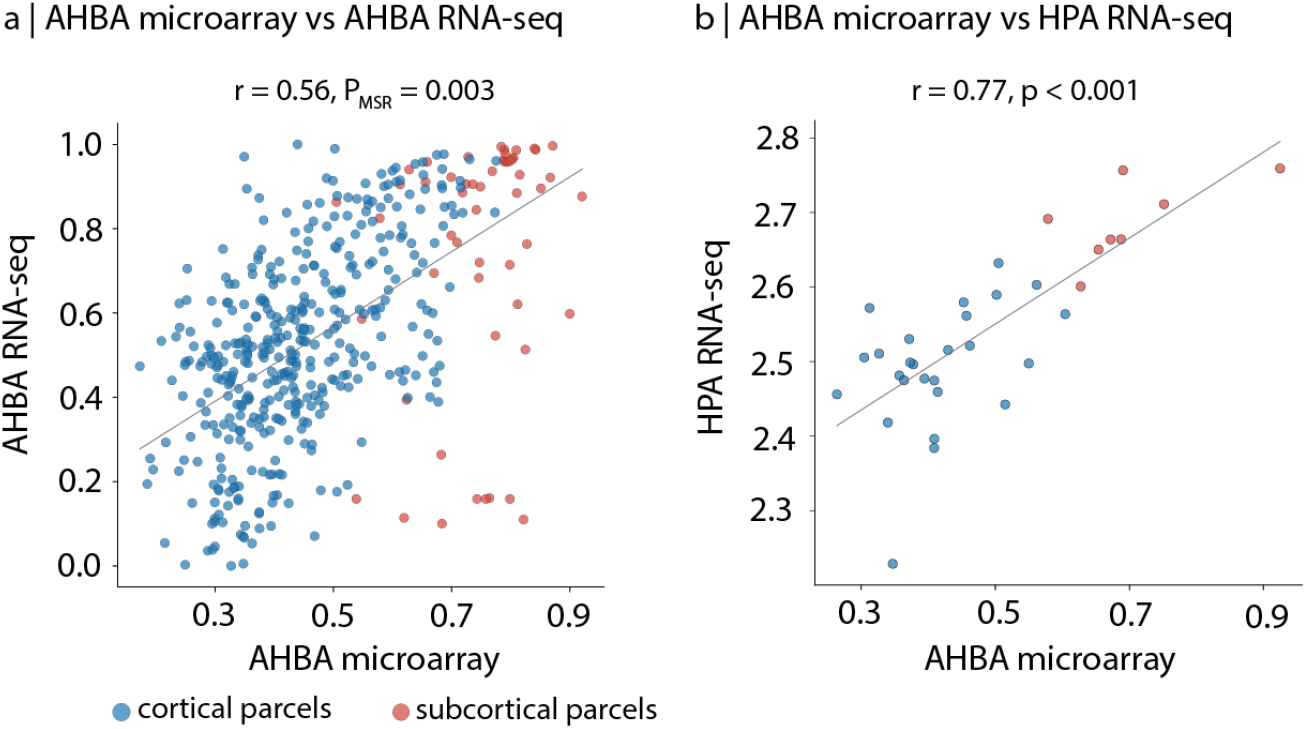
Comparisons of AQP4 expression with RNA-seq data from AHBA and the Human Protein Atlas (HPA) To assess the consistency of spatial patterns of AQP4 expression in our discovery dataset (microarray measurements from the Allen Human Brain Atlas, AHBA) [45], we systematically compare it with two independent RNA-seq datasets. (a) Parcel-wise AQP4 expression from AHBA microarray is plotted against AHBA RNA-seq estimates across cortical and subcortical parcels (Schaefer–Melbourne parcellation [107, 117]; blue, cortical parcels; red, subcortical parcels). (b) AHBA microarray AQP4 expression is parcellated according to the cortical Destrieux atlas [20] and subcortical Melbourne subcortical S1 atlas [117], then matched to RNA-seq measurements from the Human Protein Atlas [111, 119], yielding 26 cortical and 8 subcortical regions of interest. In both panels, the grey line shows the best-fitting linear regression, and the reported statistics give the correlation coefficient and associated p-value.

**Figure S2.**
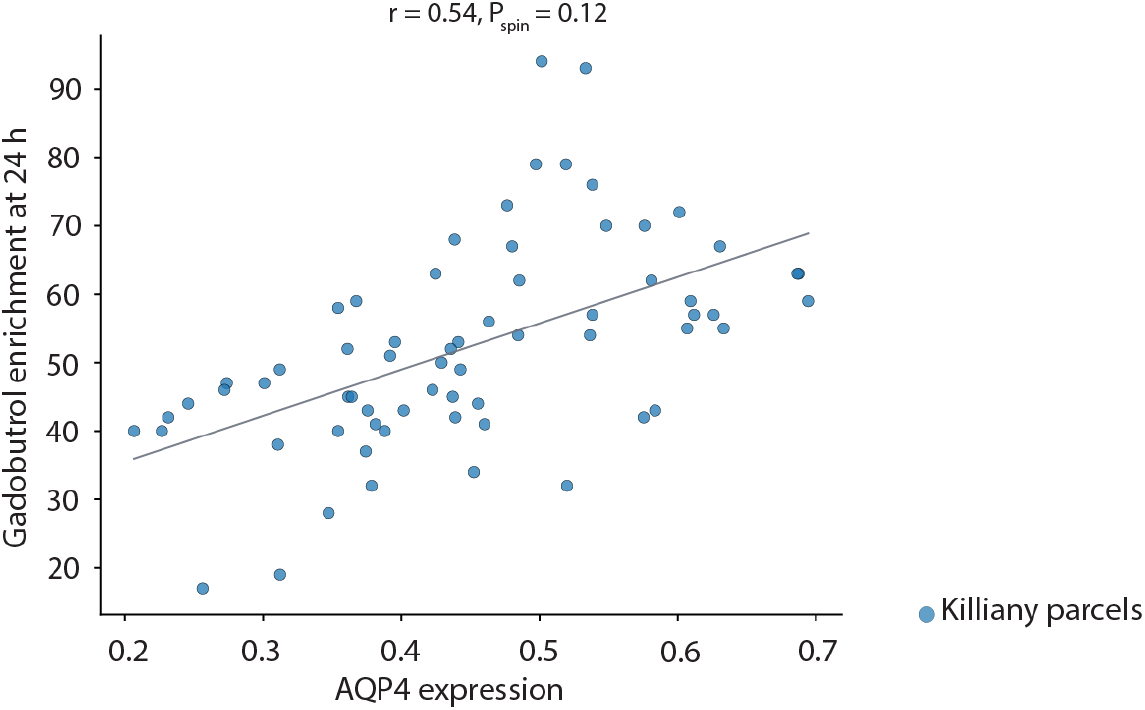
AQP4 expression correlates with delayed intrathecal CSF tracer enrichment. Scatterplot shows parcel-wise Pearson correlation between AQP4 expression estimated from AHBA microarray data [45] (x-axis) and regional enrichment of an intrathecally administered MRI tracer (gadobutrol) measured 24 hours after injection in reference participants from Ringstad et al. [101] (y-axis). Tracer enrichment values reflect the 24 hour percent change in normalized T1 signal relative to the pre-contrast baseline, quantified over FreeSurfer-derived anatomical regions [26, 27, 99]. AQP4 values are mapped to the same FreeSurfer/Desikan–Killiany parcellation used for the tracer analysis to enable direct spatial comparison. The grey line shows the best-fitting least-squares regression.

**Figure S3.**
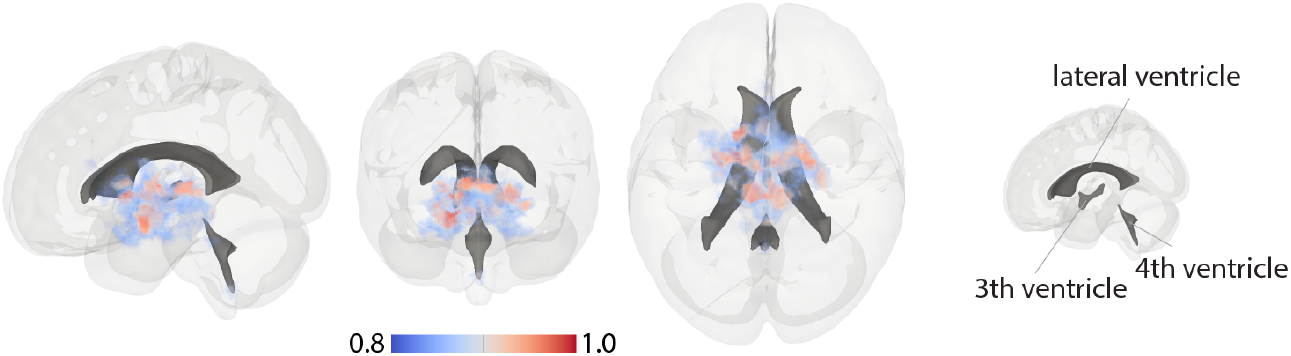
Spatial distribution of high AQP4 expression. A dense, interpolated volumetric map of AQP4 expression was generated from Allen Human Brain Atlas (AHBA) microarray data using the abagen workflow (see *Methods*) [45, 76] and rendered in MNI152NLin2009cAsym (1 mm) space. Only voxels with normalized expression values in the range 0.8– 1.0 are shown as a volume rendering (blue–red scale). Ventricular anatomy (lateral, 3rd, and 4th ventricles; FreeSurfer aseg labels 4, 43, 14, and 15) is overlaid in grey for anatomical reference, showing that high AQP4 expression tends to concentrate near ventricular spaces. Left-to-right: sagittal, coronal, and axial views.

**Figure S4.**
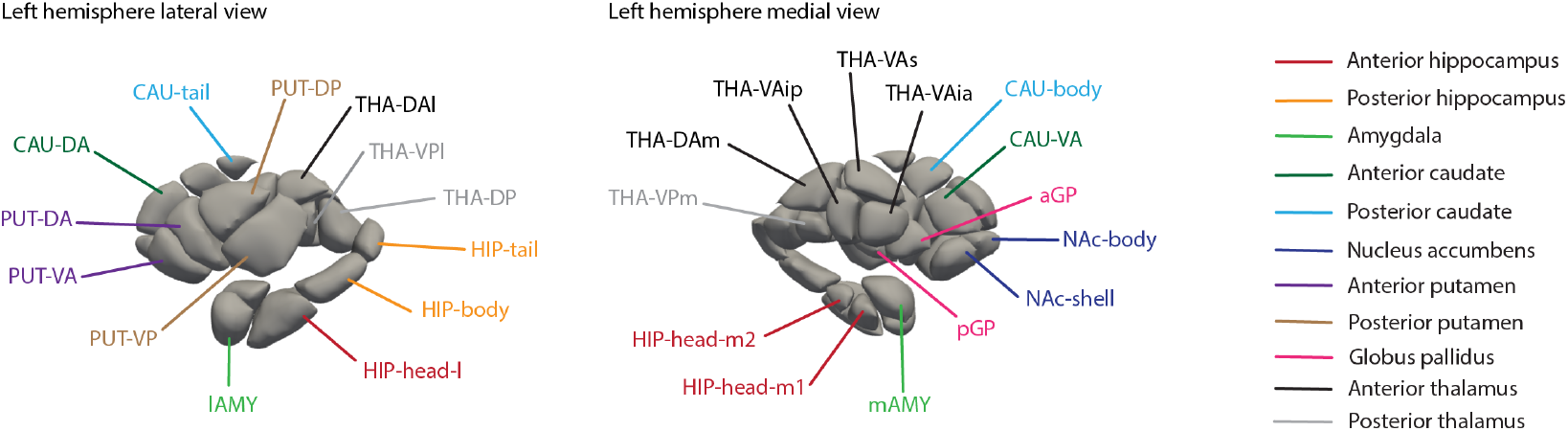
Labels for Melbourne subcortical S4 parcels. Left hemisphere lateral (left) and medial (right) views of the Melbourne subcortical S4 atlas [117] showing the parcel abbreviations. Abbreviations and full region names are listed in Table. S1.

**TABLE S1.** Nomenclature for the Melbourne subcortical S4 atlas [117] | These labels are displayed in Fig. S4.

| Abbreviation | Area's full name | Structure |
| --- | --- | --- |
| HIP-head-m1 | Hippocampus head, medial division - Subdivision 1 | Anterior hippocampus |
| HIP-head-m2 | Hippocampus head, medial division - Subdivision 2 | Anterior hippocampus |
| HIP-head-l | Hippocampus head, lateral division | Anterior hippocampus |
| HIP-body | Hippocampus body | Posterior hippocampus |
| HIP-tail | Hippocampus tail | Posterior hippocampus |
| LAMY | Lateral amygdala | Amygdala |
| mAMY | Medial amygdala | Amygdala |
| CAU-DA | Dorsoanterior caudate | Anterior caudate |
| CAU-VA | Ventroanterior caudate | Anterior caudate |
| CAU-tail | Caudate tail | Posterior caudate |
| CAU-body | Caudate body | Posterior caudate |
| NAc-shell | Nucleus accumbens, shell | Nucleus accumbens |
| NAc-body | Nucleus accumbens, body | Nucleus accumbens |
| PUT-DA | Dorsoanterior putamen | Anterior putamen |
| PUT-VA | Ventroanterior putamen | Anterior putamen |
| PUT-DP | Dorsoposterior putamen | Posterior putamen |
| PUT-VP | Ventroposterior putamen | Posterior putamen |
| aGP | Anterior globus pallidus | Globus pallidus |
| pGP | Posterior globus pallidus | Globus pallidus |
| THA-DAl | Lateral dorsoanterior thalamus | Anterior thalamus |
| THA-DAm | Medial dorsoanterior thalamus | Anterior thalamus |
| THA-VAs | Superior ventroanterior thalamus | Anterior thalamus |
| THA-VAia | Inferior ventroanterior thalamus - Anterior division | Anterior thalamus |
| THA-VAip | Inferior ventroanterior thalamus - Posterior division | Anterior thalamus |
| THA-DP | Dorsoposterior thalamus | Posterior thalamus |
| THA-VPm | Medial ventroposterior thalamus | Posterior thalamus |
| THA-VPl | Lateral ventroposterior thalamus | Posterior thalamus |

**TABLE S2.** Nomenclature for the multi-modal Glasser parcellation [33].

| Abbreviation | Area's full name | Abbreviation | Area's full name |
| --- | --- | --- | --- |
| V1 | Primary Visual Cortex | MST | Medial Superior Temporal Area |
| V6 | Sixth Visual Area | V2 | Second Visual Area |
| V3 | Third Visual Area | V4 | Fourth Visual Area |
| V8 | Eighth Visual Area | 4 | Primary Motor Cortex |
| 3b | Primary Sensory Cortex | FEF | Frontal Eye Fields |
| PEF | Premotor Eye Field | 55b | Area 55b |
| V3A | Area V3A | RSC | RetroSplenial Complex |
| POS2 | Parieto-Occipital Sulcus Area 2 | V7 | Seventh Visual Area |
| IPS1 | IntraParietal Sulcus Area 1 | FFC | Fusiform Face Complex |
| V3B | Area V3B | LO1 | Area Lateral Occipital 1 |
| LO2 | Area Lateral Occipital 2 | PIT | Posterior InferoTemporal complex |
| MT | Middle Temporal Area | A1 | Primary Auditory Cortex |
| PSL | PeriSylvian Language Area | SFL | Superior Frontal Language Area |
| PCV | PreCuneus Visual Area | STV | Superior Temporal Visual Area |
| 7Pm | Medial Area 7P | 7m | Area 7m |
| POS1 | Parieto-Occipital Sulcus Area 1 | 23d | Area 23d |
| v23ab | Area ventral 23 a+b | d23ab | Area dorsal 23 a+b |
| 31pv | Area 31p ventral | 5m | Area 5m |
| 5mv | Area 5m ventral | 23c | Area 23c |
| 5L | Area 5L | 24dd | Dorsal Area 24d |
| 24dv | Ventral Area 24d | 7AL | Lateral Area 7A |
| SCEF | Supplementary and Cingulate Eye Field | 6ma | Area 6m anterior |
| 7Am | Medial Area 7A | 7Pl | Lateral Area 7P |
| 7PC | Area 7PC | LIPv | Area Lateral IntraParietal ventral |
| VIP | Ventral IntraParietal Complex | MIP | Medial IntraParietal Area |
| 1 | Area 1 | 2 | Area 2 |
| 3a | Area 3a | 6d | Dorsal area 6 |
| 6mp | Area 6mp | 6v | Ventral Area 6 |
| p24pr | Area Posterior 24 prime | 33pr | Area 33 prime |
| a24pr | Anterior 24 prime | p32pr | Area p32 prime |
| a24 | Area a24 | d32 | Area dorsal 32 |
| 8BM | Area 8BM | p32 | Area p32 |
| 10r | Area 10r | 47m | Area 47m |
| 8Av | Area 8Av | 8Ad | Area 8Ad |
| 9m | Area 9 Middle | 8BL | Area 8B Lateral |
| 9p | Area 9 Posterior | 10d | Area 10d |
| 8C | Area 8C | 44 | Area 44 |
| 45 | Area 45 | 47l | Area 47l (47 lateral) |
| a47r | Area anterior 47r | 6r | Rostral Area 6 |
| IFJa | Area IFJa | IFJp | Area IFJp |
| IFSp | Area IFSp | IFSa | Area IFSa |
| p9-46v | Area posterior 9-46v | 46 | Area 46 |
| a9-46v | Area anterior 9-46v | 9-46d | Area 9-46d |
| 9a | Area 9 anterior | 10v | Area 10v |
| a10p | Area anterior 10p | 10pp | Polar 10p |
| 11l | Area 11l | 13l | Area 13l |
| OFC | Orbital Frontal Complex | 47s | Area 47s |
| LIPd | Area Lateral IntraParietal dorsal | 6a | Area 6 anterior |
| i6-8 | Inferior 6-8 Transitional Area | s6-8 | Superior 6-8 Transitional Area |
| 43 | Area 43 | OP4 | Area OP4/PV |
| OP1 | Area OP1/SII | OP2-3 | Area OP2-3/VS |
| 52 | Area 52 | RI | RetroInsular Cortex |
| PFcm | Area PFcm | PoI2 | Posterior Insular Area 2 |
| TA2 | Area TA2 | FOP4 | Frontal OPercular Area 4 |
| MI | Middle Insular Area | Pir | Piriform Cortex |
| AVI | Anterior Ventral Insular Area | AAIC | Anterior Agranular Insula Complex |
| FOP1 | Frontal OPercular Area 1 | FOP3 | Frontal OPercular Area 3 |
| FOP2 | Frontal OPercular Area 2 | PFt | Area PFt |
| AIP | Anterior IntraParietal Area | EC | Entorhinal Cortex |
| PreS | PreSubiculum | H | Hippocampus |
| ProS | ProStriate Area | PeEc | Perirhinal Ectorhinal Cortex |
| STGa | Area STGa | PBelt | ParaBelt Complex |
| A5 | Auditory 5 Complex | PHA1 | ParaHippocampal Area 1 |
| PHA3 | ParaHippocampal Area 3 | STSda | Area STSd anterior |
| STSdp | Area STSd posterior | STSvp | Area STSv posterior |
| TGd | Area TG dorsal | TE1a | Area TE1 anterior |
| TE1p | Area TE1 posterior | TE2a | Area TE2 anterior |
| TF | Area TF | TE2p | Area TE2 posterior |
| PHT | Area PHT | PH | Area PH |
| TPOJ1 | Area TemporoParietoOccipital Junction 1 | TPOJ2 | Area TemporoParietoOccipital Junction 2 |
| TPOJ3 | Area TemporoParietoOccipital Junction 3 | DVT | Dorsal Transitional Visual Area |
| PGp | Area PGp | IP2 | Area IntraParietal 2 |
| IP1 | Area IntraParietal 1 | IP0 | Area IntraParietal 0 |
| PFop | Area PF opercular | PF | Area PF Complex |
| PFm | Area PFm Complex | PGi | Area PGi |
| PGs | Area PGs | V6A | Area V6A |
| VMV1 | VentroMedial Visual Area 1 | VMV3 | VentroMedial Visual Area 3 |
| PHA2 | ParaHippocampal Area 2 | V4t | Area V4t |
| FST | Area FST | V3CD | Area V3CD |
| LO3 | Area Lateral Occipital 3 | VMV2 | VentroMedial Visual Area 2 |
| 31pd | Area 31pd | 31a | Area 31a |
| VVC | Ventral Visual Complex | 25 | Area 25 |
| s32 | Area s32 | pOFC | Posterior OFC Complex |
| PoI1 | Area Posterior Insular 1 | Ig | Insular Granular Complex |
| FOP5 | Area Frontal Opercular 5 | p10p | Area posterior 10p |
| p47r | Area posterior 47r | TGv | Area TG Ventral |
| MBelt | Medial Belt Complex | LBelt | Lateral Belt Complex |
| A4 | Auditory 4 Complex | STSva | Area STSv anterior |
| TE1m | Area TE1 Middle | PI | Para-Insular Area |
| a32pr | Area anterior 32 prime | p24 | Area posterior 24 |

